# Tree phyllospheres are a habitat for diverse populations of CO-oxidising bacteria

**DOI:** 10.1101/2021.03.12.435102

**Authors:** Jess L. Palmer, Sally Hilton, Emma Picot, Gary D. Bending, Hendrik Schäfer

## Abstract

**Background:** Carbon monoxide (CO) is a naturally occurring and ubiquitous trace gas in the atmosphere. As a product of combustion processes, it can reach concentrations in the mg/m^3^ range in urban areas, contributing to air pollution. Aerobic CO-degrading microorganisms have been identified previously and are thought to remove ~370 Tg of CO in soils and oceans per year. Based on the presence of genes encoding subunits of the enzyme carbon monoxide dehydrogenase in metagenomes, a large fraction of soil bacteria may have the potential for CO degradation. The activity and diversity of CO-degrading microorganisms in above ground habitats such as the phyllosphere has not been addressed, however, and their potential role in global CO cycling remains unknown.

**Results:** Monitoring of CO-degradation in leaf washes of two common British trees, *Ilex aquifolium* and *Crataegus monogyna*, demonstrated CO uptake in all samples investigated. Leaf washes of *I. aquifolium* had significantly higher CO oxidation rates than those of *C. monogyna*. A diverse range of bacterial taxa were identified as candidate CO-oxidising taxa based on high-throughput sequencing and multivariate statistical analysis of 16S rRNA amplicon data, as well as functional diversity analysis based on *coxL*, the gene encoding the large subunit of CO-dehydrogenase. Candidate CO-oxidising taxa included a range of Rhizobiales and Burkholderiales, of which the Burkholderiales OTUs were abundant colonisers of the phyllosphere at the time of sampling, as indicated by 16S rRNA gene sequencing. In addition, an estimated 12.4% of leaf OTUs in samples of this study contained *coxL* homologues, based on their predicted genomes. We also mined data of publicly available phyllosphere metagenomes for genes encoding subunits of CO-dehydrogenase which indicated that, on average, 25% of phyllosphere bacteria contained CO-dehydrogenase gene homologues. A CO-oxidising Phyllobacteriaceae strain was isolated from phyllosphere samples which contains genes encoding both CODH as well as a RuBisCO.

**Conclusions:** The phyllosphere, a vast microbial habitat, supports diverse and potentially abundant CO-oxidising bacteria. These findings identify tree phyllosphere bacteria as a potential sink for atmospheric CO and highlight the need for a more detailed assessment of phyllosphere microbial communities in the global cycle of CO.

## Introduction

The phyllosphere, defined as the above ground parts of plants, is a vast microbial habitat covering an estimated surface area of around 1 billion km^2^ [4, 5]. It is colonised by diverse microorganisms including fungi, archaea, protists, viruses and bacteria, of which bacteria are the most abundant group, with an estimated 10^6^ – 10^7^ bacterial cells per cm^2^ of leaf [6]. Phyllosphere microbial communities have taxonomically diverse bacterial communities that vary seasonally, geographically and with plant host [4, 7–10].

UV radiation, fluctuations in water availability and temperature, and scarcity of nutrients, characterise the phyllosphere as a harsh environment. Phyllosphere bacteria have been implicated in plant health, and interactions with the plant host range from symbiotic to pathogenic [11, 12]. Phyllosphere bacteria also contribute to biogeochemical cycling and degradation of pollutants, as demonstrated by the degradation of plant or atmospherically derived organic compounds such as methanol [13], phenol [14], 4-chlorophenol [15], benzene [16], polycyclic aromatic hydrocarbons (PAHs) [17], methyl chloride [18] isoprene [19, 20] and diesel [21]. In doing so, activities of phyllosphere bacteria affect atmospheric chemistry and the fate of atmospheric pollutants and may thus contribute to critical ecosystem services of the phyllosphere in mitigation of air pollution, a major global public health problem causing around 5.5 million premature deaths worldwide [22].

Carbon monoxide (CO) is a ubiquitous component of the atmosphere. It is toxic to humans and even small changes in CO concentration in urban areas can have an impact on human health, with a 1 mg/m^3^ increase in CO being associated with a 4.4% increase in cardiovascular disease hospital admissions [23]. Atmospheric CO also negatively impacts the environment by reducing concentrations of hydroxyl radicals and thus increasing the residence times of greenhouse gases such as methane [24], contributing to the increase in ground-level ozone [25] and leading to the formation of photochemical smog [26].

Environmental sources of atmospheric CO include volcanic eruptions, bushfires, photochemical reactions and photolysis of marine coloured dissolved organic matter (CDOM)[27, 28]. In addition, both live and dead plant matter have been shown to emit CO due to photochemical transformations within living leaf tissue or by UV-induced photoproduction by dead leaf tissue. This results in an estimated 50-200 Tg yr^-1^ of CO and 60-90 Tg yr^-1^ of CO being produced from live plant tissues and dead plant matter, respectively [29, 30].

The major global source of CO, however, is anthropogenic combustion processes which contribute more than half of annual CO emissions (approx. >1,200 Tg per year) [31, 32]. CO concentrations are highest in urban areas, ranging from 1.1 – 2.5 mg/m^3^ in the UK [33], compared to the atmospheric background of roughly 50-150 ppb (0.058 – 0.17 mg/m^3^) [34–36]. However, in polluted cities such as Beijing, CO concentrations as high as 17.1mg/m^3^ have been reported [37].

Most CO in the atmosphere is converted to CO2 via tropospheric hydroxylation [38], but significant amounts of CO are removed by microbial oxidation. Approximately 10-15% of the global CO flux is consumed by soil microbiota [39] which remove approximately 300 Tg of CO per year [32]; marine microbiota are estimated to oxidise approximately 70 Tg of CO per year [40], preventing the release of 85% of CO produced in the oceans into the atmosphere [41, 42]. Despite the fact that a significant amount (110-290 Tg yr^-1^) of CO is produced by living and dead plant matter, it is currently unknown whether plant-associated microorganisms play a role in the mitigation of CO from this source.

Bacteria using CO as a carbon and energy source, so-called carboxydotrophs, typically utilise high concentrations of CO (>1%), assimilating the CO2 produced via the Calvin cycle. However, carboxydotrophs have a low affinity for CO, preferring other organic substrates [32]. Other CO-oxidisers, known as carboxydovores, cannot grow at elevated CO concentrations but oxidise CO up to 1000 ppm. Carboxydovores have a high affinity for CO but cannot generate biomass when CO is the sole carbon source; they likely use CO as a supplementary energy source [32].

Aerobic CO-oxidising bacteria oxidise CO to CO2 using the enzyme carbon monoxide dehydrogenase (CODH) of which two forms, referred to as form I and form II, have been described. Both enzymes are composed of three subunits, encoded by *coxL* (large subunit), *coxM* (medium subunit), and *coxS* (small subunit) [43]. The form I *cox* operon is transcribed in the order M, S, L whereas the form II operon is transcribed in the order S, L, M. Form I CODH has been described as the ‘definitive’ form and has been well-characterised in several carboxydotrophs [44]. A form II enzyme was shown to be a functional CODH in *Bradyrhizobium japonicum* USDA 110 [45] and *Kyrpidia spormannii* [46], but the inability to oxidise CO by a range of bacteria harbouring exclusively form II CODH [47] suggests that some aspects of form II CODH function or regulation remain unknown. Carboxydotrophs and carboxydovores are taxonomically diverse, including members of the Proteobacteria, Firmicutes, Actinobacteria, Chloroflexi and Bacteroidetes identified in marine [47–49], soil [48, 50, 51] and rhizosphere environments [45, 48]. The presence of *coxL* genes in a wide range of environments and in representatives of uncultivated clades of bacteria indicates that the diversity of CO-oxidising bacteria is greater than those currently identified by cultivation [49, 52–58]. CO oxidation is known to enhance the long-term survival of some bacteria, supporting their persistence in deprived or changeable environments [59] and recent research indicates that CO oxidation may be a more generalist function than previously assumed, with 56% of soil bacteria predicted to be capable of CO oxidation [60]. It is therefore possible that CO oxidation may be a widespread function in a range of microbial environments.

CO emissions from plant matter and anthropogenic sources such as vehicle exhaust fumes likely provides a constant input of CO to the leaf environment, suggesting that CO could be a nutrient exploited by bacteria in the nutrient-limited habitat of the leaf surface. Here we assessed the potential of phyllosphere microbial communities to degrade CO and identified CO-degrading microorganisms using a combination of cultivation-dependent and cultivation-independent approaches. We show that tree phyllosphere samples are capable of CO oxidation and that some samples oxidised CO even when no other organic carbon sources were provided. The phyllosphere communities of *Ilex aquifolium* (holly) and *Crataegus monogyna* (hawthorn) contained diverse populations of CO degrading bacteria based on high-resolution microbial community analysis coupled to multivariate statistical analyses and functional marker diversity of *coxL*. A Phyllobacteriaceae strain able to oxidise CO was also isolated from phyllosphere samples. Overall, these results identify tree phyllospheres as a habitat for CO-oxidising bacteria and suggest that phyllosphere microorganisms contribute to global cycling of CO.

## Materials and Methods

### Sample collection and processing

Two species of common trees in Britain, hawthorn (*Crataegus monogyna*) and holly (*Ilex aquifolium*) were investigated, providing samples from both a deciduous and evergreen tree species. Trees were sampled on 6/10/2017. Tree branches were sampled in Coventry, United Kingdom, with hawthorn trees sampled at both a woodland (sample code ‘HtW’) (52.376248, −1.551322) and roadside (sample code ‘HtR’)(52.394371, −1.554182) location whereas holly trees were sampled at the woodland site (sample code ‘HlW’) (52.376248, −1.556771) only. At each site, four trees were selected and whole twigs were removed at random from 1.5-2m high and were placed into sterile, polyethylene zip-lock bags to be transported to the lab where they were processed within 24 hours.

Phyllosphere communities were collected by adapting a leaf washing method by Atamna-Ismaeel et al., [61]. In summary, 5 g of leaves per sample were weighed into sterile 250 ml Erlenmeyer flasks, to which 150 ml sterile phosphate-buffered saline (PBS) buffer (137 mM NaCl, 2.7 mM KCl, 10 mM Na2HPO4, 1.8 mM KH2PO4, pH 7.4) was added. Microorganisms and other particulates were then dislodged from the leaf surface by sonication for 2 minutes in a water bath, followed by 10 seconds of vortexing and a 5-minute resting period. The vortex and resting stages were repeated 6 times to thoroughly dislodge leaf surface material into the buffer. Leaf wash was filtered through 0.22 μm pore size membrane filters (Millipore, USA). Leaf wash filters were cut in half and either immediately used to determine their potential for CO oxidation or immediately frozen and stored at −80°C for future DNA extraction.

### Assessment of leaf wash CO oxidation potential and cultivation of phyllosphere microbial communities enriched with CO

Sterile mineral salts medium was prepared for use in CO degradation experiments and enrichment of CO-degrading microorganisms as described by Meyer and Schlegel [44]. Analytical grade chemicals were dissolved in Milli-Q water before sterilisation by autoclaving at 121°C for 15 minutes. To encourage the heterotrophic oxidation of CO, 0.25 g/L of yeast extract and 1 mL of filter sterilised vitamin solution [62] was added to the mineral salts medium. Solid media for isolation of CO-oxidising bacteria were prepared accordingly with addition of 1.5% (w/v) BactoAgar (Becton Dickinson, USA). Liquid media (25 mL) added to 125 mL serum vials were inoculated by adding half of a leaf wash filter per vial. Vials were sealed with a butyl plug, crimp-sealed, and CO was injected into the headspace to a concentration of approximately 800 ppm. Sterile controls were prepared without inoculum and no-substrate controls were prepared containing the microbial inoculum but without addition of CO. Enrichment cultures and controls were incubated in the dark in a shaking incubator at 22°C and 100 rpm.

Headspace CO concentrations were monitored every 2-4 days using an Agilent 6890N gas chromatograph (GC; Agilent Technologies, California, US) equipped with a nickel catalyst methaniser (Agilent Technologies, California, US) allowing detection of CO after its reduction to methane using a flame ionisation detector. The system was fitted with a 1.5 m long 80/100 Porapak Q column (inner diameter 2.1 mm; Sigma-Aldrich, US) held at 250°C using nitrogen (50 mL/min) used as the carrier gas. A 100 μl headspace sample was injected into the apparatus and peak readings for CO were taken. CO headspace concentrations in the headspace of enrichment cultures were calculated based on a calibration derived from injection of samples of known CO concentrations. CO concentrations between 0 (ambient air) and 50 ppm could not be differentiated and so <50 ppm was defined as the detectable limit.

Once CO concentrations were below the detection threshold, sub-cultures were prepared for both the enrichment sample and its corresponding control sample. Enrichment sub-cultures were prepared and monitored as above, using 2 mL of liquid culture from the corresponding sample as the microbial inoculum added to 23 mL of fresh mineral salts medium (plus yeast extract). This was repeated twice upon the degradation of CO for a total of three sub-culture cycles per sample. Once the final sub-culture had degraded the CO, 15 mL of liquid culture was extracted and centrifuged at 4500 × g for 20 minutes to form a pellet. The supernatant was removed and the pellet was flash frozen in liquid nitrogen before storing at −80°C for future DNA extraction.

Once individual samples had degraded CO on their final sub-culture, they and their corresponding no-substrate control culture were also subcultured into mineral salts media without addition of yeast extract so that CO was the sole carbon source in order to investigate the occurrence of autotrophic degradation of CO. These ‘autotrophic’ subcultures were prepared by centrifugation of 15 mL of the final ‘heterotrophic’ sub-culture at 4500 × g for 10 minutes, discarding the supernatant and resuspending the pellet in 2 mL of autotrophic CO. This was repeated three times before the washed pellet resuspension was added as an inoculum to 23 mL of mineral salts media without yeast extract. Degradation of CO was monitored and cultures were re-spiked with CO to bring the headspace concentration back to 800 ppm upon depletion. A total of three re-spikes were monitored and upon degradation of the final CO pulse, 15 mL of liquid culture was processed as described for heterotrophic CO enrichment cultures above for isolation of colonies and DNA extraction.

### DNA extraction

DNA from CO enrichment cultures and no-substrate controls was extracted using the FastDNA^™^ Spin Kit for Soil (MP Bio Science Ltd., Derby, UK). The required amount of sodium phosphate buffer and MT buffer was added to the enrichment culture pellet in order to resuspend the pellet before being transferred to the Lysing Matrix E tube and proceeding with the protocol as per the manufacturer’s instructions. At the final step, DNA was eluted using 50 μL of elution buffer. Aliquots of DNA extracts were assessed for quantity and quality by gel electrophoresis using 0.8% (w/v) agarose gels and a Qubit® 2.0 Fluorometer (Invitrogen, USA), using high-sensitivity reagents.

### 16S rRNA PCR and MiSeq library preparation

High-throughput sequencing was used to investigate the diversity of 16S rRNA genes in CO enrichment culture samples, corresponding no-substrate controls and original leaf wash samples. For each sample, 20 ng of DNA was used as template to amplify 292bp of 16S rRNA genes using the 515f (5′-GTGCCAGCMGCCGCGGTAA-3′) and the 806r (5′-GGACTACHVGGGTATCTAAT-3′) primer pair [63] with Illumina Nextera Transposase Adapters attached to the 5′ end. PCRs were performed in 25 μL reaction volumes and contained 13 μL Q5^®^ Hot Start High-Fidelity 2× Master Mix (New England Biolabs, US), 0.5 μM of each primer and 0.4 μL of 25 mg/mL BSA. The reaction conditions were as follows: an initial denaturing step of 98°C for 30 s followed by 25 cycles of 98°C for 10 s, 50°C for 15 s and 72°C for 20 s and a final extension of 72°C for 10 mins. PCR products were then purified using the AmpliClean^™^ Magnetic Bead PCR Clean-Up Kit (NIMAGEN, Nijmegen, Netherlands), which was carried out as per the manufacturer’s instructions.

Purified PCR products were given unique dual indexes at the 5’ end using the Nextera XT Index Kit v2 index primers (Illumina, USA). To attach the index primers, a PCR mixture was made consisting of 13 μL Q5^®^ Hot Start High-Fidelity 2× Master Mix, 1 μM Index Primer 1, 1 μM Index Primer 2 and 4 μl of purified PCR product, with the addition of sterile, nuclease-free dH_2_O for a final reaction volume of 26 μL per sample. The unique indexes were added by amplifying PCR products with the following reaction conditions: 95°C for 3 mins, followed by 8 cycles of 98°C for 20 s, 55°C for 15 s, and 72°C for 15 s, then a final elongation step of 72°C for 5 mins. Samples were normalised using the SequalPrep^TM^ Normalisation Plate Kit (Invitrogen, USA) as per the manufacturer’s instructions. All samples were pooled and adjusted to 4 nM before being submitted to the Genomics Facility at the University of Warwick, Coventry, United Kingdom for sequencing on the Illumina Miseq platform.

### *coxL* functional gene PCR amplification and construction of clone libraries

Amplification of form I *coxL* genes using DNA from CO enrichment cultures was initially attempted using the OMPf and O/Br *coxL* primer pair [48]. As this was not successful, this primer pair was reviewed and modified. Form I *coxL* sequences were obtained from the Integrated Microbial Genome and Metagenome (IMG) database [64] using the BLASTx function with a *Burkholderia fungorum* form I *coxL* sequence (AY307914.1) as a query. Five-hundred extra form I *coxL* genes were obtained in addition to the nine form I *coxL* genes used for the design of the original OMPf O/Br primer pair which were then aligned using ARB software [65] and a neighbour-joining tree was constructed. Primer sequences were reviewed with focus on *coxL* genes from terrestrial origins. An alignment was made with a total of 437 *coxL* sequences, the diversity of which is displayed in Figure S1. Mismatches were identified and modifications made to increase the redundancy of the primers in an attempt to improve upon the range of bacterial species targeted by the primers. Modifications made are shown in Table S1.

The newly modified forward (5′- GGCGGNTTYGGNAAYAARGT) and reverse (5′- YTCDATDATCATNGGRTTDAT) primer pair was used to amplify definitive (form I) *coxL* genes from samples. PCRs were prepared containing 1x KAPA Taq Buffer A (includes 1.5 mM MgCl2) (Kapa Biosystems, USA), dNTP nucleotide mix (0.2 mM each), DMSO (2% v/v), BSA (0.05% w/v), forward and reverse primer (0.4 μM each), KAPA Taq DNA polymerase (2 U) (KAPABIOSYSTEMS) and DNA template (2-20 ng). The reaction was made up to a 50 μL reaction volume with sterile nuclease-free dH_2_O. Positive controls were included which contained DNA template from *Ruegeria pomeroyi* DSS-3, a known CO oxidiser. Negative controls did not contain any DNA template. Reaction conditions were as follows: initial denaturation at 94°C for 5 min, followed by 35 cycles of a 30 s denaturation at 94°C, 30 s annealing at 56°C and 1:15 min extension at 72°C, with a final extension of 7 min at 72°C. PCR products were visualised with GelRed (Biotium, USA) gel electrophoresis and gel-purified. PCR products were sequenced by Sanger sequencing either by GATC Biotech GmbH (Konstanz, Germany) or Eurofins Genomics (Ebersberg, Germany).

The diversity of *coxL* genes in CO enrichment culture samples was assessed by cloning purified PCR products using the TOPO TA cloning kit and TOP10 competent cells (Invitrogen, USA) as per the manufacturer’s instructions. Transformants were used as template for PCR amplification of *coxL* using the modified primer pair. Products were purified and sequenced as described previously.

### Isolation, identification and genome sequencing of CO-oxidising bacteria

Microbial culture (100 μl) sampled from heterotrophic or autotrophic enrichment vials after the final sub-culture was serially diluted with mineral salts media [44]. Dilutions were spread plated onto mineral salts plates (containing 0.25 g/L yeast extract) and LB plates and incubated at 22°C in the dark. Single colonies were selected at random and screened by PCR for presence of *coxL* gene as described previously. If products of the correct size were amplified, the corresponding colony was then re-suspended into mineral salts medium with 800 ppm CO and monitored for CO degradation by GC.

For taxonomic identification of isolates, the 16S rRNA gene was amplified as described above for *coxL* PCR but using the 27f (5′- AGAGTTTGATCMTGGCTCAG) and 1492r (5′- TACGGYTACCTTGTTACGACTT) primer pair [66] and using 1μl of cell suspension from isolates that had first been heated at 100°C for 5 min. Reaction conditions were as follows: initial denaturation at 94°C for 5 min, followed by 35 cycles of a 30 s denaturation at 94°C, 30 s annealing at 55.5°C and 1:15 min extension at 72°C, with a final extension of 7 min at 72°C. PCR products were visualised, purified and sequenced as described previously.

For whole genome sequencing of the CO-oxidising isolate, a lawn of cells was grown on an LB plate before harvesting of the cells which were sent to MicrobesNG (Birmingham, UK) where DNA was extracted and sequenced using the Illumina HiSeq platform (250bp paired-end protocol). Adapters were trimmed and assembled by MicrobesNG using Trimmomatic [67] and SPAdes version 3.7 [68], respectively. Contigs were then submitted to RAST [69] for annotation and analysis of CO oxidation gene clusters.

### Bioinformatic and statistical analyses

For the processing of MiSeq data, low-quality sequences were removed from the ends of the pre-demultiplexed reads using Trimmomatic version 0.36 [67]. USEARCH [70] was then used for the majority of downstream analyses. Paired ends were joined, quality filtering was applied to a 280 bp minimum, primers were removed, and samples were dereplicated to remove any singletons. OTUs were clustered to a 97% identity threshold which also removed the majority of chimeras. Remaining chimeras were removed using the GOLD database [71] for 16S rRNA amplicons. Reads were then mapped back to the OTU database for generation of OTU tables.

QIIME [72] was used along with the Greengenes [73] reference database to assign taxonomy to 16S rRNA gene amplicons. OTU tables were generated and any OTUs identified as mitochondria or chloroplasts were removed. Samples were rarefied to 9000 reads per sample defined by the sample with the lowest number of reads.

Non-metric multi-dimensional scaling (MDS) analysis of Bray Curtis similarity of bacterial community structure were conducted using the VEGAN package in R [74]. Analysis of Similarity (ANOSIM) tests were used to determine how dissimilar bacterial communities were between groups using PRIMER v6 software [75]. Candidate CO-oxidising OTUs were identified by a combination of Similarity Percentages (SIMPER) analysis using PRIMER v6 software [75], to find those which drove the most dissimilarity between CO enrichment cultures and controls, and identification of OTUs which showed a significant log2fold increase in CO enrichment cultures compared to controls, which was done using the DESeq2 package [76] and visualised using the EnhancedVolcano package [77] in RStudio.

Significant differences in CO degradation times between sample types were calculated using Wilcoxon tests in R studio (version 1.1.447, RStudio Inc., US) to calculate P values.

All sequence traces were checked for quality using 4Peaks (Nucleobytes.com, Aalsmeer, The Netherlands) and high-quality sequence was used as a query for BLASTn (nucleotide BLAST for 16S rRNA sequences) or BLASTx (translated nucleotides protein BLAST for functional genes) searches [78]. Sequences were aligned with their top hits from the BLAST database using MUSCLE [79] in MEGA7 [2]. Phylogenetic trees were constructed using the neighbour-joining algorithm [1] with a bootstrap test applied [3] with 500 replicates in MEGA7.

The PICRUSt2 [80] package was used to predict the functional capability of the original leaf wash community to oxidise CO, based on 16S rRNA amplicon data. 10649 OTUs were used in the analysis, of which 9701 (91%) had a closely related genome available on the IMG database. From the predicted genomes of the OTUs and the relative abundance of each OTU (based on 16S rRNA data), the predicted copy number of *coxL* (KEGG K03520) was then compared between leaf wash sample types.

### Mining of CODH homologues from phyllosphere metagenomes

In addition to predictions made by PICRUSt2 of the abundance of *coxL* genes from samples in this study, the metagenomics-rapid annotation using subsystems technology (MG-RAST) database [81] was used to determine how abundant CODH genes are in available phyllosphere metagenomes. Abundance data for protein homologues (identified by the MG-RAST pipeline) of CODH subunits (CoxL, CoxM and CoxS), methanol dehydrogenase subunits (MxaF and MxaI) and three housekeeping genes *(recA, rho* and *rpoA*) were extracted from all publicly available phyllosphere metagenomes which included clover, soybean, *Arabidopsis thaliana* [82], rice [8] and 48 neotropical tree samples [83]. The abundances of Cox and methanol dehydrogenase homologues were then normalised against the three housekeeping genes to determine the relative abundance of the target functional genes in the bacterial communities. Attempts to align a sub-set (77 AA sequences) of clover metagenome CoxL amino acid sequences in order to confirm that sequences were CoxL homologues was unsuccessful due to the short length (~60-90 AA) of partial sequences of CoxL homologues. However, BLASTp searches indicated that the majority had high (>70%) AA sequence identity to other CoxL or xanthine dehydrogenase (XDH) homologues. In addition, when the same subset (77 AA sequences) of CoxL homologues were aligned with both characterised CoxL [48] and xanthine dehydrogenase large subunits [84], the majority (68/77) had a higher sequence identity with CoxL than with xanthine dehydrogenase sequences.

## Results

### Leaf wash communities are capable of CO oxidation

Incubation of leaf wash samples from all three sample types, woodland hawthorn (HtW), roadside hawthorn (HtR) and woodland holly (HlW) in mineral medium amended with yeast extract resulted in the degradation of 800 ppm CO (Figure 1). On average, it took 33 days for CO to be degraded beyond the detectable limit (<50 ppm) by all phyllosphere samples (n=12), although there was variation in different replicate samples (range of 17-52 days). This demonstrated that leaf wash samples, and hence phyllosphere microbial communities, contained viable microorganisms capable of CO degradation.

**Figure 1:**
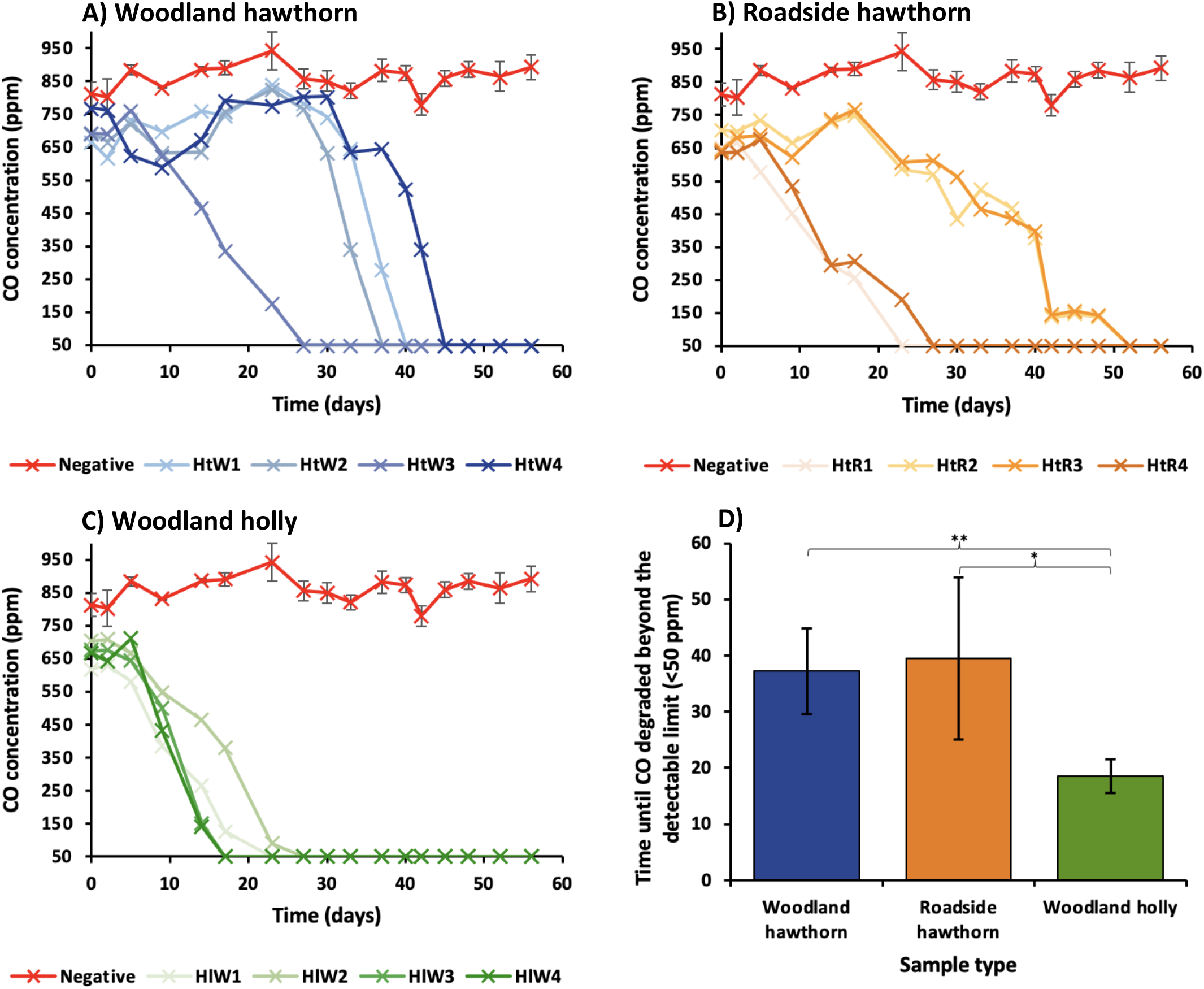
Degradation of CO by holly and hawthorn leaf wash communities cultivated in media with added yeast extract. Enrichment culture medium contained 800_ppm CO with 0.25_g/L yeast extract. **A)** Woodland hawthorn (HtW) samples. **B)** Roadside hawthorn (HtR) samples. **C)** Woodland holly (HlW) samples. **D)** Average time taken for CO degradation between sample types (** = p-value of <0.001, * = p-value of <0.05, error bars indicate standard deviation). Negative controls (n=4) did not contain leaf wash as an inoculum. CO concentrations displayed in this figure are those of enrichment cultures where original leaf wash samples were used as an inoculum. CO oxidation data from subsequent sub-cultures is not shown.

The length of lag phases between samples also varied, with 5/8 hawthorn samples (HtW and HtR) showing long lag phases of between 17-30 days whereas some hawthorn samples (3/8) and all holly samples had short lag phases of 2-5 days before degradation of CO began. Once degradation of CO began, woodland samples degraded CO faster (average 45 ppm and 39 ppm CO per day for HtW and HlW samples, respectively), than the roadside hawthorn samples (24 ppm CO per day).

Woodland holly (HlW) samples degraded the CO the fastest (average 18.5 days), followed by woodland hawthorn (HtW) samples (average 37.25 days). Roadside hawthorn (HtR) samples were the slowest (average of 43 days). There was no significant difference between CO degradation rates of HtW and HtR samples (P value = 0.49). However, HlW samples degraded CO significantly faster than HtW (P value = 0.0037) and HtR (P value = 0.013) samples.

Once each sample had been sub-cultured three times upon CO degradation, the samples were inoculated into minimal media with CO but without additional carbon source to potentially indicate the presence of autotrophic CO oxidisers. Only half of all samples subcultured in this way continued to fully degrade three spikes of 800ppm CO, suggesting that some of the enrichments contained carboxydotrophic (autotrophic) CO oxidisers in addition to carboxydovores (Figure S2). It is noteworthy that those cultures which continued to degrade CO generally showed increasing rates of CO degradation which might indicate increased abundance of autotrophic CO oxidisers.

### Candidate CO oxidisers in the phyllosphere include rare and abundant community members

Following complete CO degradation of three consecutive sub-cultures or re-spikes, the diversity of the enrichment cultures and no-substrate controls was assessed by high-throughput sequencing of 16S rRNA genes. Analyses of the bacterial community composition outlined below revealed how leaf surface communities responded to CO during the incubations and identified candidate CO oxidisers.

ANOSIM statistics showed that CO enrichment cultures were weakly but significantly dissimilar to their corresponding no-substrate controls, irrespective of whether they contained additional yeast extract as carbon source or not (R= 0.31, P = 0.001 and R=0.24, P= 0.02, respectively), as shown in Figure 2. Differences in bacterial community composition between CO enrichment cultures and controls at the OTU level are shown in Figures S3 and S4. No significant dissimilarities were found in the community composition of cultures inoculated with leaf wash from different tree species or tree locations (data not shown).

**Figure 2:**
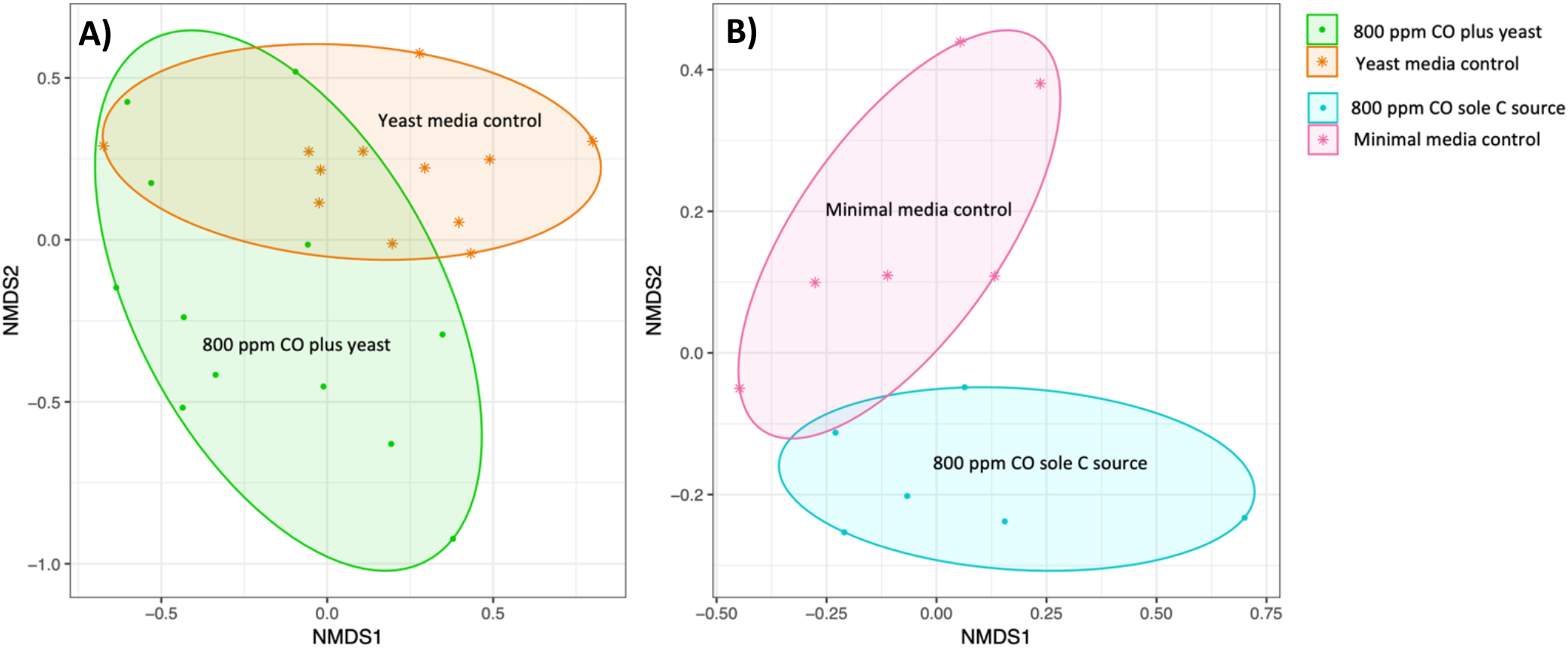
MDS ordination of variation between bacterial OTUs of CO enrichment culture samples and no-substrate control incubations. MDS analyses were derived from Bray-Curtis similarity matrices constructed with relative read abundance data of OTUs present in each sample. **A)** Variation between leaf wash samples cultivated with or without 800 ppm CO with the addition of another carbon source (yeast extract). **B)** Variation between subcultures from leaf wash samples cultivated with 800 ppm CO plus yeast extract which were then able to oxidise 800 ppm CO as a sole carbon source and no-substrate controls.

SIMPER analysis identified those OTUs that drove the dissimilarity between the leaf wash cultures enriched with CO and no-substrate controls, thus highlighting candidate CO oxidisers. For this purpose, we initially considered candidate CO oxidisers as those OTUs that were amongst the top ten OTUs driving dissimilarity between CO and no-substrate controls, and which showed >1% increase in relative abundance in the CO enrichment cultures versus the no-substrate controls (Table 1). The majority of these OTUs were present at low relative abundances in the unenriched leaf wash samples. However, three candidate CO oxidiser OTUs (OTU9, OTU36 and OTU10) together comprised >0.5 – 5.5% of total leaf wash community reads, on average. Strikingly, OTU10 (Comamonadaceae) had high relative read abundance (range 2.1-10.1%) in all unenriched leaf wash communities suggesting it could be an abundant coloniser of the phyllosphere.

**Table 1:**
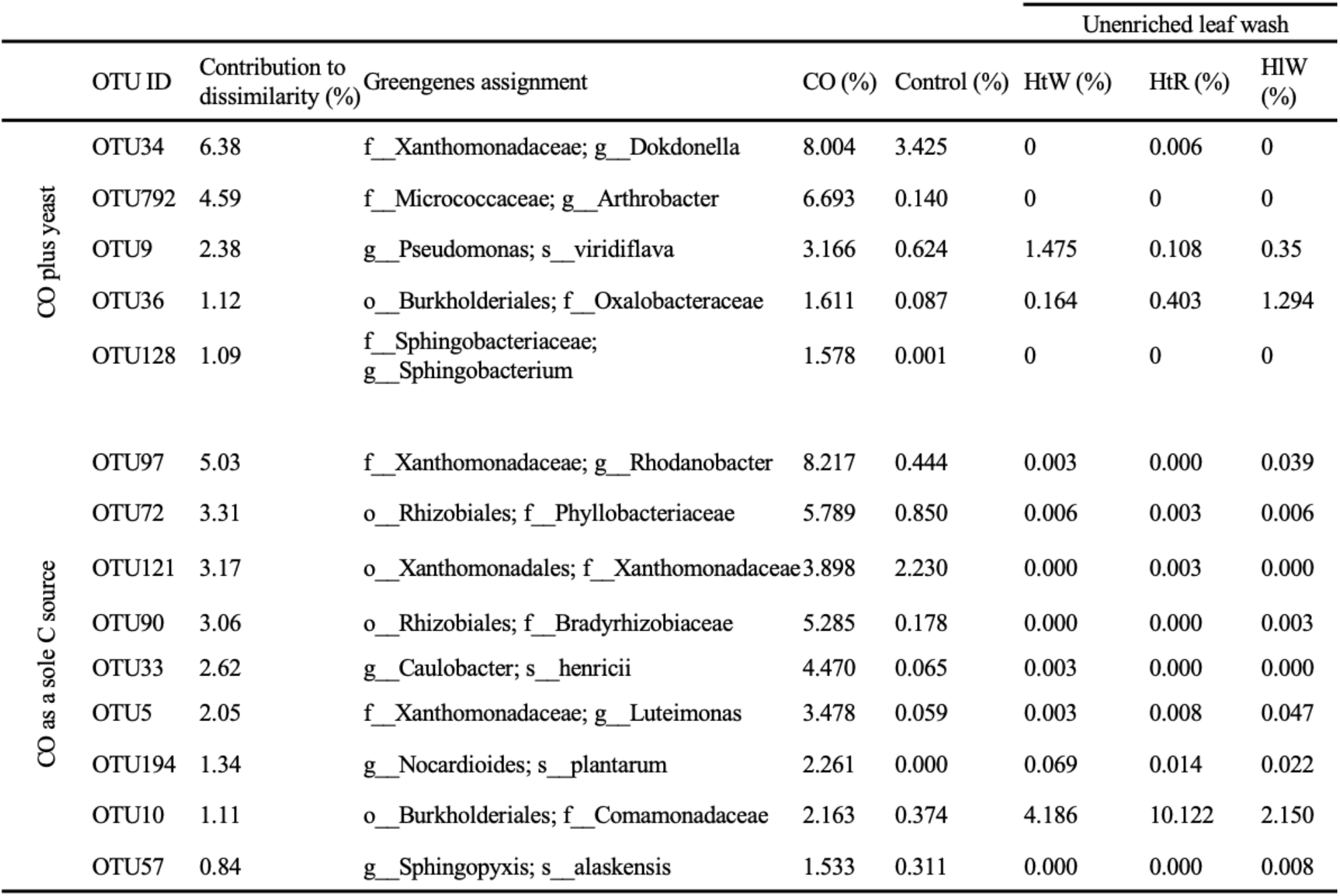
Candidate CO-oxidising OTUs based on SIMPER analyses. Values represent relative abundance. Where Greengenes assignments are indicated, o_ = order, f_ = family, g_ = genus and s_ = species. Where values for unenriched leaf wash samples are indicated, HtW = woodland hawthorn samples, HtR = roadside hawthorn samples and HlW = woodland holly samples.

|  | OTU ID | Contribution to dissimilarity (%) | Greengenes assignment | CO (%) | Control (%) | Unenriched leaf wash |  |  |
| --- | --- | --- | --- | --- | --- | --- | --- | --- |
|  |  |  |  |  |  | HtW (%) | HtR (%) | HIW (%) |
| CO plus yeast | OTU34 | 6.38 | f__Xanthomonadaceae; g__Dokdonella | 8.004 | 3.425 | 0 | 0.006 | 0 |
|  | OTU792 | 4.59 | f__Micrococcaceae; g__Arthrobacter | 6.693 | 0.140 | 0 | 0 | 0 |
|  | OTU9 | 2.38 | g__Pseudomonas; s__viridiflava | 3.166 | 0.624 | 1.475 | 0.108 | 0.35 |
|  | OTU36 | 1.12 | o__Burkholderiales; f__Oxalobacteraceae | 1.611 | 0.087 | 0.164 | 0.403 | 1.294 |
|  | OTU128 | 1.09 | f__Sphingobacteriaceae; g__Sphingobacterium | 1.578 | 0.001 | 0 | 0 | 0 |
| CO as a sole C source | OTU97 | 5.03 | f__Xanthomonadaceae; g__Rhodanobacter | 8.217 | 0.444 | 0.003 | 0.000 | 0.039 |
|  | OTU72 | 3.31 | o__Rhizobiales; f__Phyllobacteriaceae | 5.789 | 0.850 | 0.006 | 0.003 | 0.006 |
|  | OTU121 | 3.17 | o__Xanthomonadales; f__Xanthomonadaceae | 3.898 | 2.230 | 0.000 | 0.003 | 0.000 |
|  | OTU90 | 3.06 | o__Rhizobiales; f__Bradyrhizobiaceae | 5.285 | 0.178 | 0.000 | 0.000 | 0.003 |
|  | OTU33 | 2.62 | g__Caulobacter; s__henricii | 4.470 | 0.065 | 0.003 | 0.000 | 0.000 |
|  | OTU5 | 2.05 | f__Xanthomonadaceae; g__Luteimonas | 3.478 | 0.059 | 0.003 | 0.008 | 0.047 |
|  | OTU194 | 1.34 | g__Nocardioides; s__plantarum | 2.261 | 0.000 | 0.069 | 0.014 | 0.022 |
|  | OTU10 | 1.11 | o__Burkholderiales; f__Comamonadaceae | 2.163 | 0.374 | 4.186 | 10.122 | 2.150 |
|  | OTU57 | 0.84 | g__Sphingopyxis; s__alaskensis | 1.533 | 0.311 | 0.000 | 0.000 | 0.008 |

Using the Deseq2 package, we further considered as potential candidate CO oxidisers all of those OTUs which showed a log2-fold increase of >1 and a *P* value of <0.05 in cultures enriched with CO versus no-substrate controls (Table 2 and Figure S5). Four of these OTUs (OTU792, OTU90, OTU33 and OTU5) had high read abundances (range 3.5 to 6.7%) in the CO enrichment cultures and had also been identified by SIMPER analysis. Therefore, in addition to contributing to the most dissimilarity between CO enrichment culture and control groups, these OTUs also showed a significant log2-fold change. All other candidate CO oxidiser OTUs identified using this approach were not identified by SIMPER analysis and were present in CO enrichment cultures at low relative read abundances which were generally <1% (except OTU87, 2.1%). Similar to candidate CO-oxidiser OTUs identified by SIMPER analysis, most of the OTUs identified in Table 2 had low relative read abundance (<0.1%) in the unenriched leaf wash samples. Noteworthy exceptions include OTU20 and OTU87, which comprised 0.26% and 0.37% of unenriched leaf wash reads on average, respectively. Additionally, OTU13, which showed a significant log2-fold change in the ‘autotrophic’ CO enrichment culture was abundant in the leaf wash samples, comprising 2% on average of reads of woodland hawthorn samples, 1.4% of roadside hawthorn samples and 4.3% of woodland holly samples. Taken together, these analyses suggest that candidate CO oxidisers in the phyllosphere microbiomes included both rare and abundant community members.

**Table 2:**
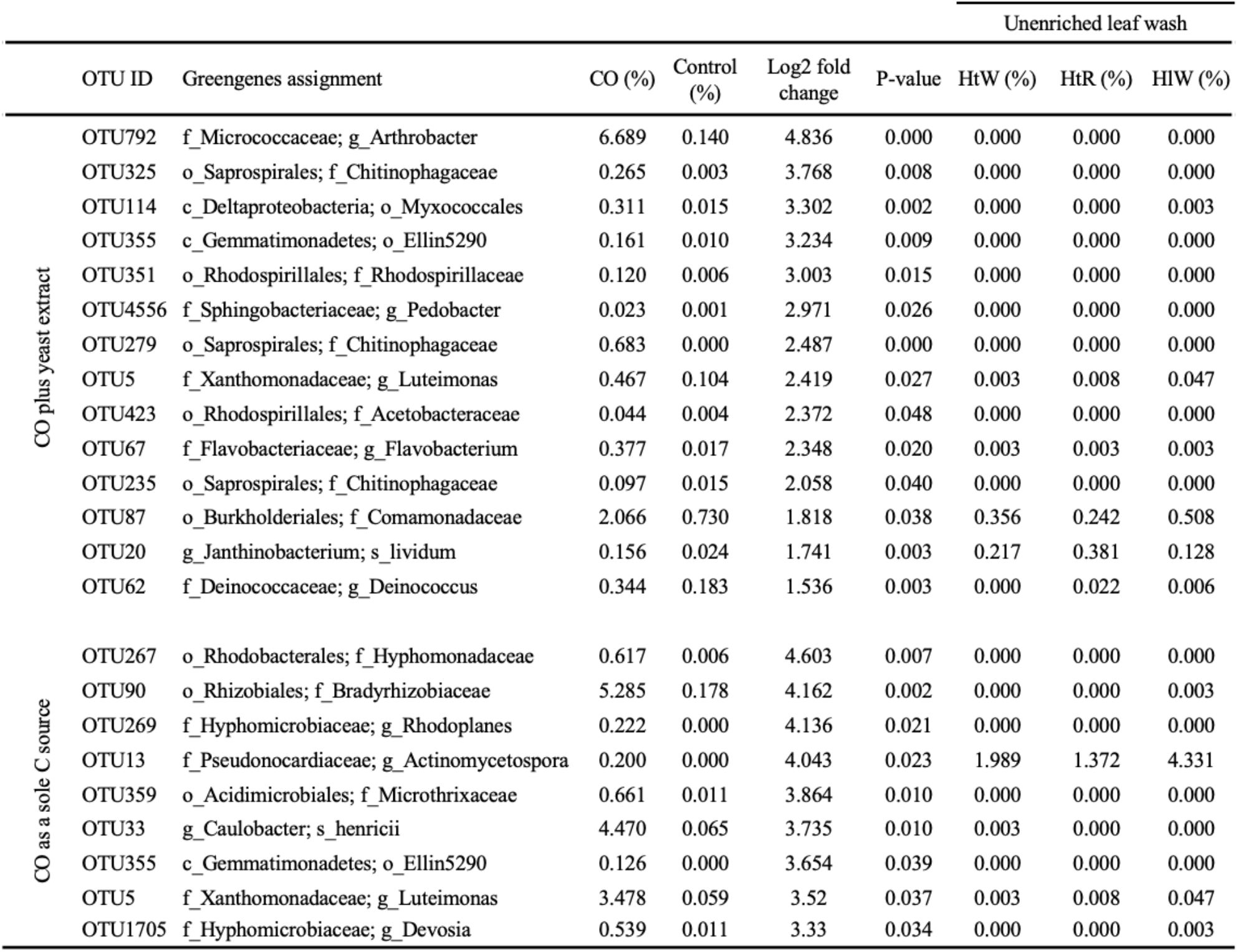
Candidate CO-oxidising OTUs indicated by those which showed the most significant log2 fold change between CO enrichment cultures and controls. Values represent relative abundance. Where greengenes assignments are indicated, o_ = order, f_ = family, g_ = genus and s_ = species. Where values for unenriched leaf wash samples are indicated, HtW = woodland hawthorn samples, HtR = roadside hawthorn samples and HlW = woodland holly samples.

### Candidate CO oxidisers in the phyllosphere include diverse bacterial taxa

The candidate CO oxidising OTUs identified above (Table 1 and 2) included a diverse range of bacteria belonging to five different phyla (Figure 3). The majority of candidate CO oxidisers were Proteobacteria (19/31), followed by Bacteroidetes (6/31), Actinobacteria (4/31), Deinococcus-Thermus (1/31) and Gemmatimonadetes (1/31). Three orders of bacteria contained the highest number of candidate CO-oxidising OTUs, four OTUs each were members of Burkholderiales, Xanthomonadales and Rhizobiales.

**Figure 3:**
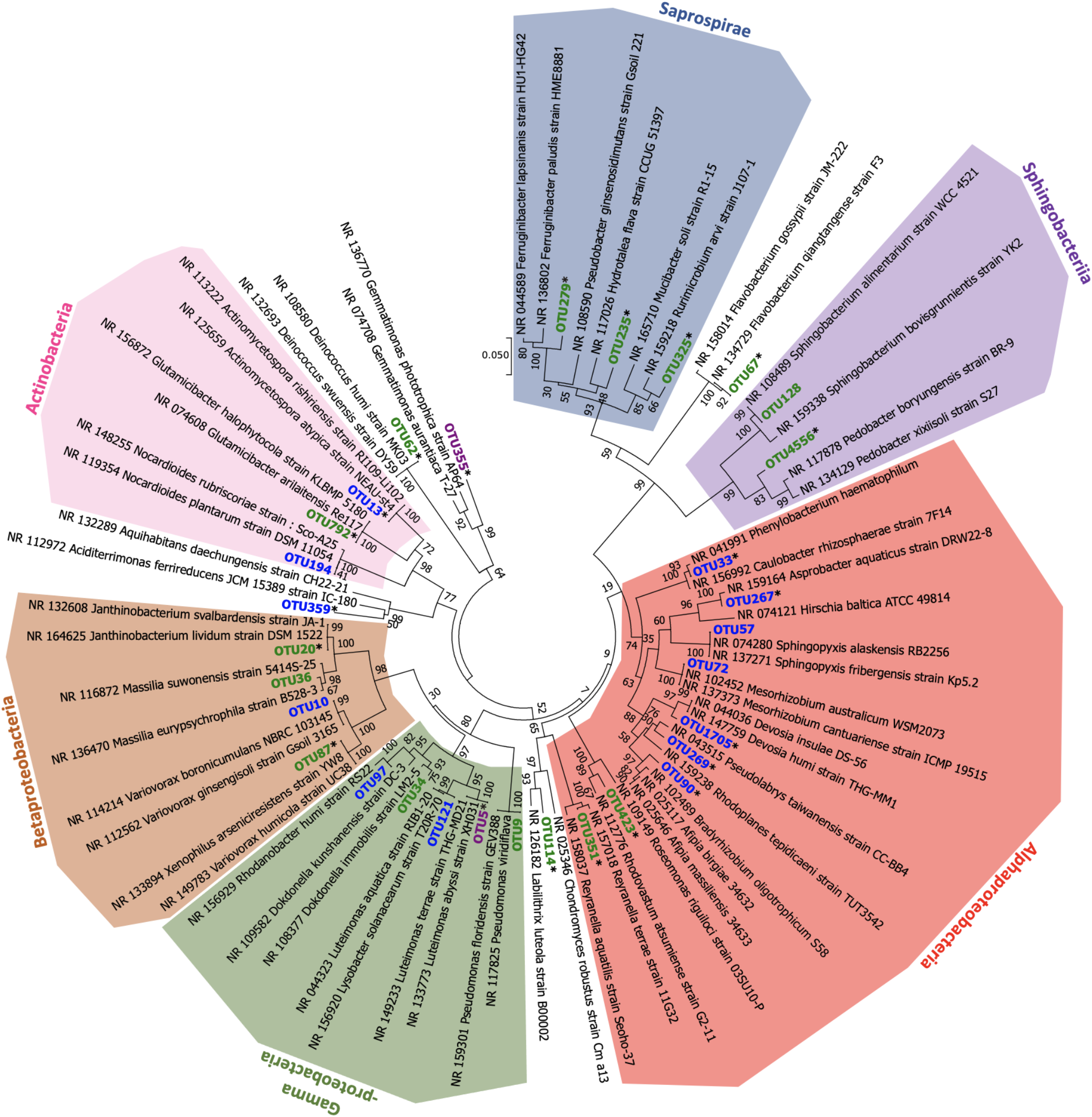
16S rRNA phylogeny of candidate CO-oxidising bacteria from phyllosphere samples. OTUs were identified as potential CO-oxidising species from SIMPER and log2 fold change analyses between leaf wash samples cultivated with CO and substrate controls without CO. OTUs indicated in green are those which were enriched in cultures where yeast extract was provided as a carbon source in addition to CO. OTUs indicated in blue are those which were enriched in cultures where CO was the sole carbon source. OTUs indicated in purple are those which were enriched in cultures where yeast extract was provided in addition to CO as well as CO as a sole carbon source. OTUs indicated with an Asterisk (*) are those which had a significant (p-value = <0.05) log fold change between CO enrichment cultures and the control group without CO. Phylogeny of OTUs and BLAST hits was inferred using the Neighbor-Joining method [1] of nucleotide sequences in MEGA7 [2]. The percentage of replicate trees in which the associated taxa clustered together in the bootstrap [3] test (500 replicates) are shown next to the branches. The scale bar denotes the number of nucleotide differences per site.

There were three groups of OTUs which clustered within their treatment group (CO plus yeast extract [Fig. 3, labelled in green] or CO only [Fig.3, labelled in blue]). The four Rhizobiales OTUs (OTU90, OTU269, OTU1705 and OTU72) were all enriched on CO when it was the sole carbon source, whereas the three Saprospirales (OTU279, OTU235 and OTU325) and the three Bacteroidetes (OTU67, OTU128, OTU4556) were identified in CO plus yeast extract incubations. Two OTUs were identified in both types of enrichments (Fig. 3, labelled purple). This analysis suggests that the phylogenetic diversity of candidate CO oxidising bacteria in the studied tree phyllospheres was high and that these might represent both carboxydotrophs and carboxydovores.

### Identification of CO-oxidising bacteria based on *cox* gene functional markers

Analysis of *coxL* diversity within CO enrichment cultures was achieved by PCR amplification, cloning and sequencing of *coxL* using three CO-enriched samples (one per sample type) with added yeast extract. Ten transformant colonies containing an insert were used in subsequent *coxL* PCRs and sent for sanger sequencing, resulting in 28/30 high-quality sequences (Figure 4).

**Figure 4:**
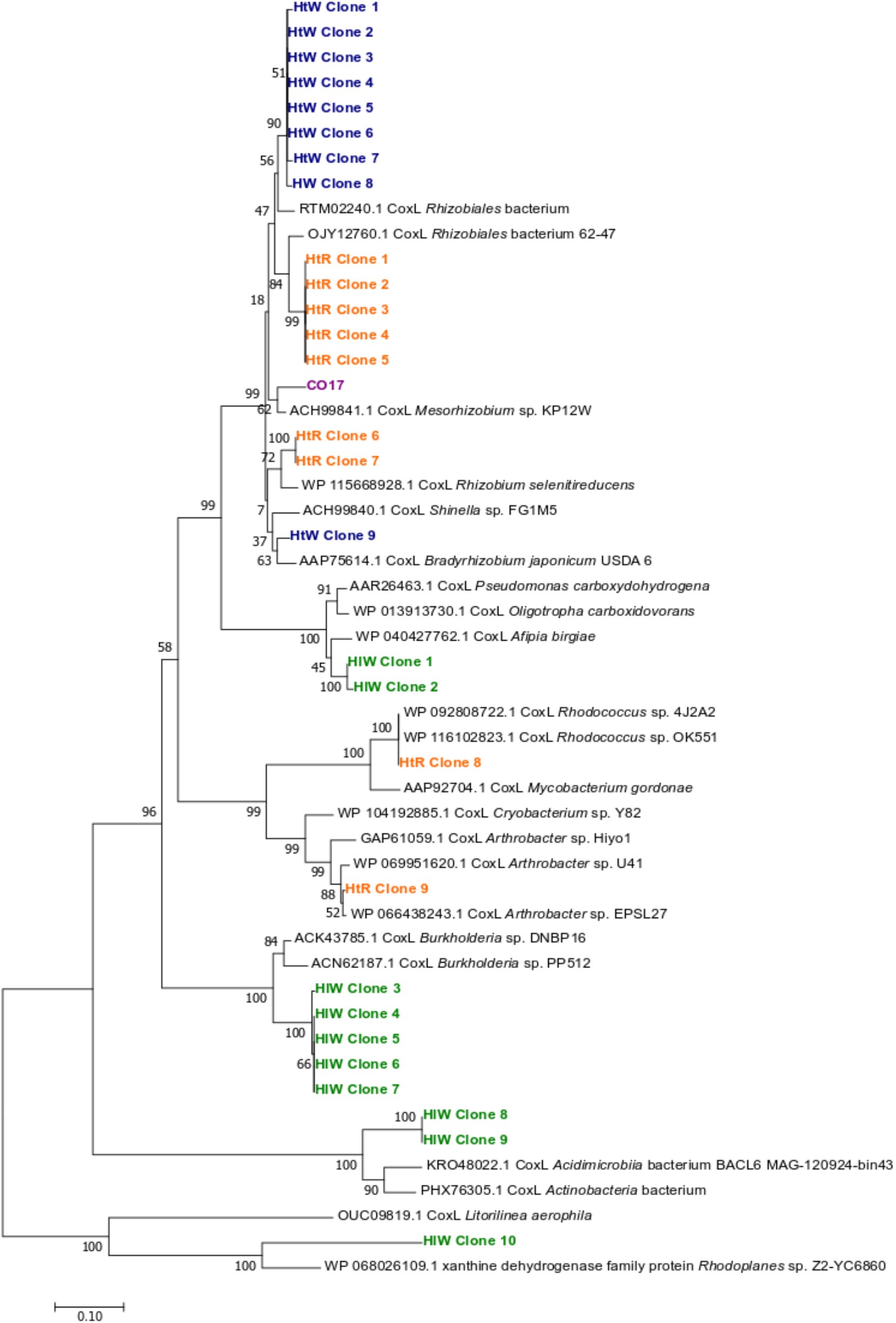
Phylogenetic tree of *coxL* PCR product sequences from CO-enrichment culture clones. Phylogeny of sequences and BLAST hits were inferred using the Neighbor-Joining method [1] of amino acid sequences in MEGA7 [2]. The percentage of replicate trees in which the associated taxa clustered together in the bootstrap [3] test (500 replicates) are shown next to the branches. Accession numbers are indicated before sequence names. HtW = woodland hawthorn samples (blue), HtR = roadside hawthorn samples (orange), HlW = woodland holly samples (green). For comparison, the CoxL amino acid sequence from isolate CO17 (purple) was also included. The scale bar denotes the number of amino acid differences per site.

Altogether, nine different clades of *coxL* sequences were identified. The majority of sequences obtained from woodland hawthorn (HtW) enrichments (8/9) were highly similar, being most closely related to *coxL* genes from a Rhizobiales bacterium and one sequence having the highest similarity to a *Bradyrhizobium coxL* gene. Four different clades of *coxL* originated from roadside hawthorn (HtR) samples, two of which were most similar to Rhizobiales, e.g. *Rhizobium selenitireducens* (2/9) and ‘Rhizobiales bacterium 62-47′ (5/9). A further two *coxL* types detected in the HtR sample were most closely related to *coxL* from *Rhodococcus* and *Arthrobacter* species, respectively. The majority of woodland holly (HlW) *coxL* sequences (5/10) were most similar to *Burkholderia coxL* genes. The remaining sequences were most similar to *coxL* from *Afipia* (2/10) or an *Acidimicrobiia* species (2/10). One of the sequences (HlW clone 10) had the highest similarity to a Xanthine dehydrogenase rather than a *coxL* gene. Thus, despite the limited depth of sequencing, the analysis of cloned *coxL* amplicons revealed a relatively high diversity of phyllosphere bacteria with the potential to degrade CO in the enrichment cultures. It is noteworthy that the closest relatives of *coxL* identified were in good agreement with some candidate CO oxidisers identified based on analysis of the 16S rRNA sequencing data, i.e. members of the Rhizobiales, Burkholderiales, and Actinobacteria. None of the cloned *coxL* sequences were identical to the *coxL* gene of CO17, a CO-oxidising phyllosphere isolate (discussed below).

PICRUSt2 [85] was used to predict how many OTUs in the leaf wash samples had a close relative which had a *coxL* homologue, the gene encoding the large subunit of CO-dehydrogenase, in their genome sequence. 948 OTUs were removed from the sample group as they did not have a close relative in the database, leaving 9701 (91%) OTUs in the analysis. Of these, PICRUSt2 analysis predicted that 12.4% (1202) of the leaf wash OTUs could have a *coxL* gene. In addition, the predicted copy number of *coxL* genes differed between some leaf wash sample types. On average, woodland holly samples contained the highest copy number of predicted *coxL* genes (5401) followed by woodland hawthorn (4867), with roadside hawthorn showing the lowest predicted *coxL* copy number (3755). While there was no significant difference (*P* = <0.05) between the hawthorn (woodland and roadside) sample types, or the woodland sample types (hawthorn and holly), woodland holly samples had a significantly higher predicted *coxL* copy number than roadside hawthorn samples (P = 0.03).

Furthermore, the relative abundance of *cox* gene homologues (normalised with three single-copy housekeeping genes) within phyllosphere metagenomes available on the MG-RAST database (clover, soybean, *Arabidopsis thaliana*, rice and neotropical trees) was determined using functional abundance data available on MG-RAST [81]. As displayed in Figure 5, the relative abundance of Cox gene homologues ranges between 21.7-55.3% for *coxL*, 10.4-27.3% for *coxM* and 9.3-28.1% for *coxS* genes. By comparison, the relative abundance of homologues of genes involved in methanol metabolism, *mxaF* and *mxaI* were between 0.5-25.9% and 0-9.8%, respectively. Overall, the average relative abundance of CODH genes (*coxL, coxM* and *coxS*) (25%) was higher than methanol dehydrogenase genes (7.5%) in the phyllosphere metagenomes which were analysed.

**Figure 5:**
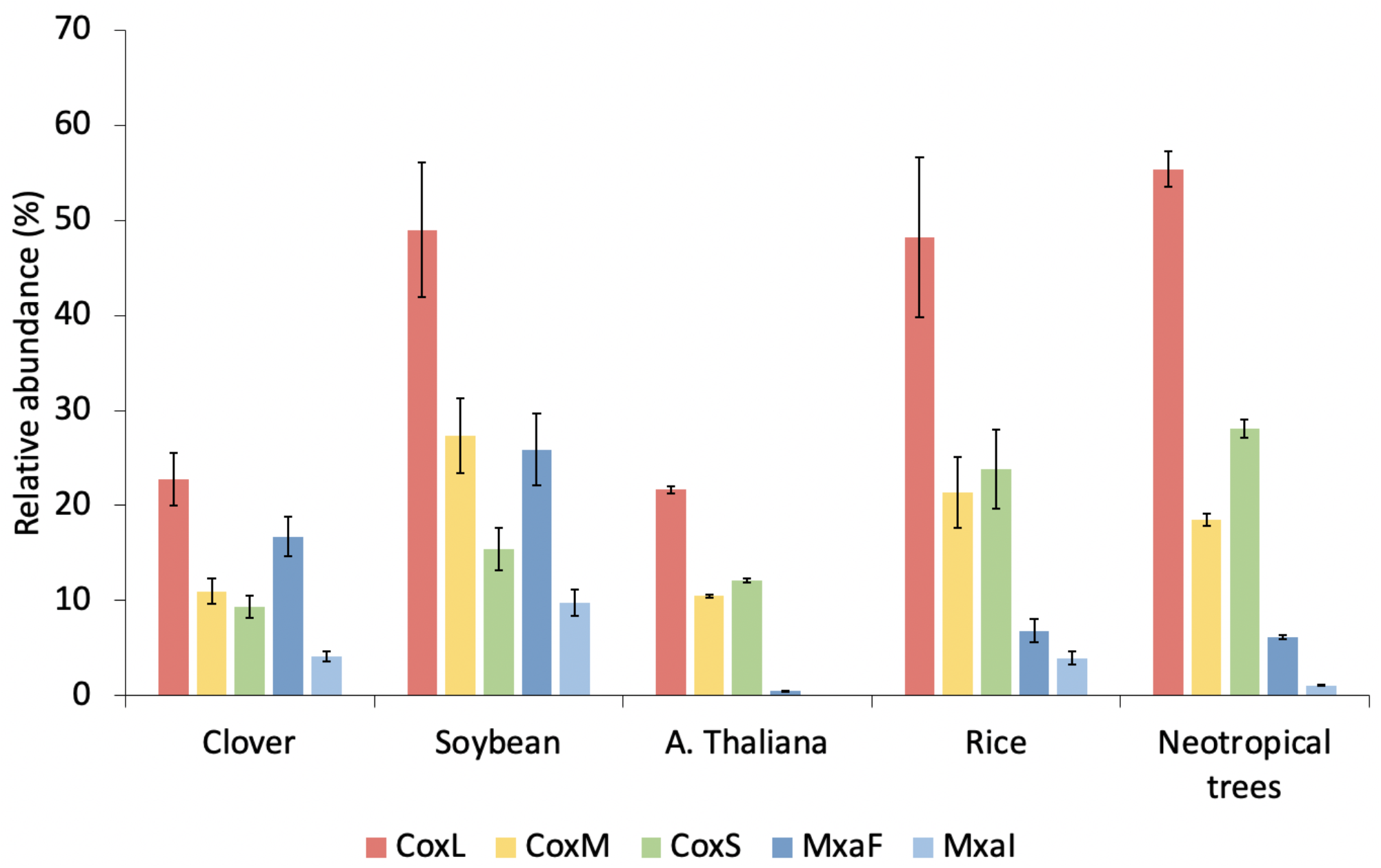
Abundance of *cox* gene homologues in phyllosphere metagenomes available on MG-RAST. Abundance of the large *(coxL*, red), medium *(coxM*, yellow) and small *(coxS*, green) subunits of CODH are shown in comparison with methanol dehydrogenase subunits *(mxaF*, dark blue and *mxaI*, light blue), a common phyllosphere function. All functional genes were normalised using three single-copy housekeeping genes: RecA, Rho and RpoA.

### A Phyllobacteriaceae isolate, strain CO17, isolated from phyllosphere samples is capable of CO oxidation

Over 20 isolates were obtained from CO enrichment cultures and screened for their ability to degrade CO, and the presence of a *coxL* gene. Only one of these isolates, designated CO17, was confirmed to oxidise CO. CO17 oxidised 800 ppm CO in the presence of yeast extract as an additional carbon source and while CO17 also oxidised 800 ppm of CO without an additional carbon source, autotrophic growth on CO as a sole carbon source could not be confirmed. Sequencing of the 16S rRNA gene identified isolate CO17 as closely related to *Mesorhizobium zhangyense*, with a 99.7% 16S rRNA sequence identity (Figure 6A). The 16S rRNA gene of isolate CO17 is distinct from that of OTU72 (a *Mesorhizobium* species identified in MiSeq analysis as a candidate CO oxidiser), sharing only 95% sequence identity. A draft genome sequence was obtained for isolate CO17 consisting of an assembly size of 7,199,351 bp (number of contigs 132, N50: 165,070; GC content 61.1%). The genome assembly of strain CO17 has operons encoding both form I and form II CODH, the arrangements of which are typical of *cox* genes in other CO-oxidising bacterial species (Figure 6B). In addition, the genome of CO17 contained homologues of both the small (*cbbS*) and large (*cbbL*) RubisCO subunits. The *cbbL* homologue of CO17 clustered within *cbbL* genes which encode the definitive form I red-like RubisCO proteins, as shown in Figure S6. This indicates that CO17 may be able to assimilate of CO2, the oxidation product of CO in autotrophic CO oxidisers.

**Figure 6:**
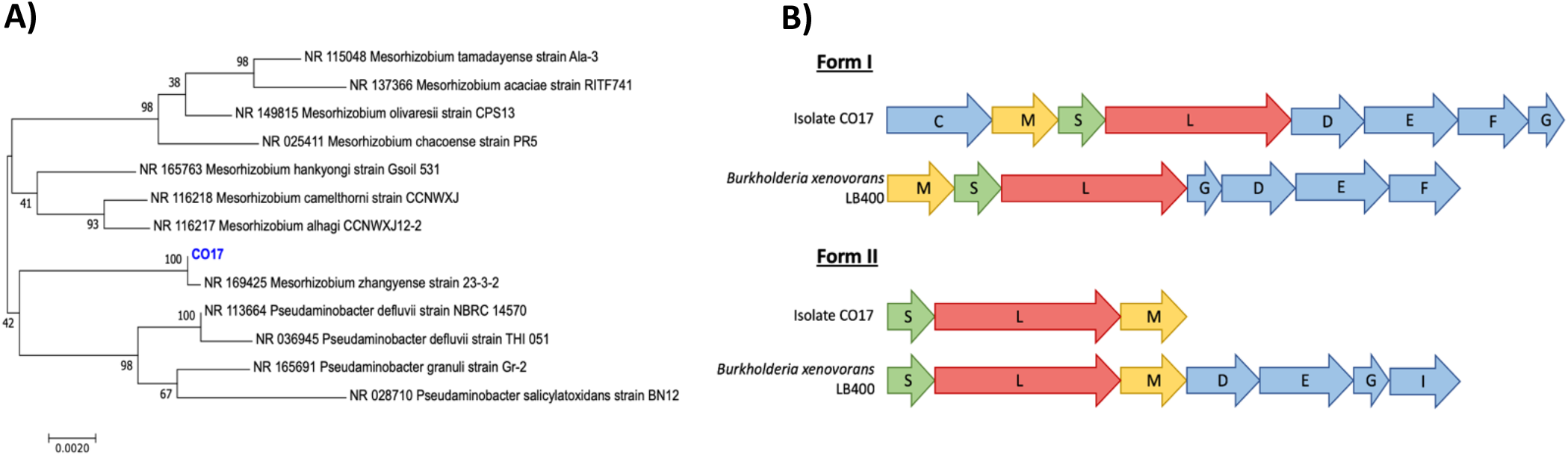
Characterisation of CO-oxidising isolate CO17. **A)** 16S rRNA phylogeny of isolate CO17. Phylogeny of isolate and BLAST hits was inferred using the Neighbor-Joining method [1] of nucleotide sequences in MEGA7 [2]. The percentage of replicate trees in which the associated taxa clustered together in the bootstrap [3] test (500 replicates) are shown next to the branches. Accession numbers are indicated before genus names. The scale bar denotes the number of nucleotide differences per site. **B)** Arrangement of form I and form II *cox* genes in isolate CO17 and *Burkholderia xenovorans* LB400. The genes encoding the small (*coxS*), medium (*coxM*) and large (*coxL*) subunits are indicated in green, yellow and red, respectively. Accessory genes are indicated in blue.

## Discussion

Although CO oxidisers have been identified previously in a range of environments, their presence in the phyllosphere has only been speculative [32]. This work provides the first evidence that there are microorganisms in the phyllosphere which are capable of CO oxidation, with all leaf wash samples in this study oxidising 800 ppm CO when cultivated with an additional carbon source. Furthermore, there may be differences in the CO-oxidising potential of different phyllosphere communities. Holly phyllosphere samples oxidised 800 ppm of CO significantly faster than hawthorn samples whereas there was no significant difference between the time taken to oxidise CO between hawthorn tree samples taken from the woodland versus the roadside. It is therefore possible that tree species has a greater impact on the ability of phyllosphere communities to oxidise CO than location. This may be due to the evergreen holly trees having a more established phyllosphere community than the hawthorn trees at the time of sampling or it may be that differences in leaf physiology between the tree species may form selective pressures on the phyllosphere community, thus altering the community composition and functional capabilities of the phyllosphere. PICRUSt data supported the idea that holly phyllosphere communities may have a higher capacity for CO oxidation, with holly samples containing the highest predicted *coxL* gene abundance. These initial data support further work that should include a wider range of tree species in order to investigate potential correlations of tree species and CO-oxidising potential of the phyllosphere.

When CO enriched leaf wash cultures were sub-cultured into media where CO was a sole carbon source, only 50% of samples were able to consistently oxidise CO. This could indicate that microorganisms capable of autotrophic CO oxidation (carboxydotrophs) are not as abundant as heterotrophic CO oxidisers (carboxydovores). This is not surprising as atmospheric CO would be available to phyllosphere microorganisms at low concentrations (~1-2 ppm in the UK) and therefore may better support microbial populations as a supplementary carbon source. Recent findings by Cordero et al. [59] showed that CODH expression is upregulated in response to starvation and, due to the abundance of CODH in soil and marine samples, they predicted that CO supported the persistence of aerobic heterotrophic bacteria in nutrient limited environments. Thus, results from this study indicating that heterotrophic CO oxidation may be more common than autotrophic CO oxidation in phyllosphere communities, the CO survival-based model of CO oxidation proposed by Cordero et al. [59] could also be an ecological adaptation enhancing survival in the phyllosphere, which is known to be deprived of energy sources.

The presence of CO in leaf wash cultures significantly impacted the bacterial community and although some of the dissimilarity between CO enrichment cultures and no-substrate controls may be due to toxic impacts of CO on the bacterial community, multiple bacterial species were identified which were potentially enriched on CO. Three candidate CO-oxidising OTUs where phylogenetically similar to genera which contain species proven to oxidise CO [44, 45, 48, 86]. These are OTU9 (a *Pseudomonas* species), OTU72 (a *Mesorhizobium* species) and OTU90 (similar to *Bradyrhizobium* species). The majority of candidate CO oxidisers therefore present species for which CO oxidation would be a novel observation for the genus they belong to. Notable examples include an *Actinomycetospora* species (OTU13), a *Phenylobacterium* species (OTU33), a *Janthinobacterium* species (OTU20) and a *Luteimonas* species (OTU5). All of which were significantly enriched in the presence of CO, indicating that CO oxidation is a likely function of these species. In addition, candidate CO-oxidising OTUs were identified within classes of bacteria which are not yet documented to contain CO-oxidising species. Species within the Acidimicrobia, Deinococci, Deltaproteobacteria, Flavobacteriia, Gemmatimonadetes, Saprospirae and Sphingobacteriia classes were identified as being potential CO oxidisers, indicating that the diversity of bacteria capable of CO oxidation may be higher than currently realised. Analysis of the functional diversity of CO oxidisers based on the *coxL* gene provided further support that some of these candidates are indeed capable of CO oxidation. Four of the candidates identified by high throughput sequencing of 16S rRNA genes were Rhizobiales species and the majority of the *coxL* sequences were most similar to those of Rhizobiales species, with five different clades of *coxL* likely belonging to this order. It is therefore likely that these candidates (OTU90, OTU269, OTU1705 and OTU72) are capable of CO oxidation. In addition, four species of Burkholderiales (OTU87, OTU10, OTU36 and OTU20) were identified in the rRNA based analyses. Interestingly, all of these OTUs were abundant in the phyllosphere at the time of sampling, making up 0.12-10% of total reads in the original leaf wash samples. A cluster of *coxL* homologues sequenced from the clone library were most similar to, but distinct from, *coxL* genes from a *Burkholderia* species. It is possible that this *coxL* gene is from one of the Burkholderiales candidates. In a parallel study investigating the degradation of 4-nitrophenol by phyllosphere microbiota (Palmer et al., unpublished), the metagenome assembled genomes (MAGs) of two species of Burkholderiales: an *Actinomycetospora* species and a *Variovorax* species believed to be OTU13 and OTU10, respectively, were obtained. These draft genomes contained the form II arrangement of *cox* genes, providing further evidence that some abundant phyllosphere colonisers are likely to be capable of CO oxidation.

Mining of CODH sequences from phyllosphere metagenomes on MG-RAST further indicate that CODH genes may be abundant within the phyllosphere microbial community. Numerous CODH homologues were detected in all the phyllosphere metagenomes available. Furthermore, when normalised with three housekeeping genes, it is estimated that up to 55% of the phyllosphere microbial community contained *coxL* gene homologues, or an average of 25% contained *cox* genes *(coxL, coxM* and *coxS*), whereas only up to 25.9% contained mxa*F* gene homologues, or an average of 7.5% from both *mxaF* and *mxaI*. Methanol metabolism is a well-known function of the phyllosphere due to the high availability of plant-produced methanol as a substrate on the leaf surface [4, 8, 82]. Therefore, these preliminary functional analyses provide an initial indication that CO oxidation has the potential to be an equally important and widespread function of phyllosphere microbiota. The estimated abundance of bacteria capable of CO oxidation in the phyllosphere in this study compliments recent estimates that 56% of soil bacteria are capable of CO oxidation [60]. It is possible that the phyllosphere and soil may harbour similar communities of CO-oxidising bacteria as these biomes seed one another by aerial dispersal, leaf fall, precipitation runoff and plant seedling inoculation. Due to the short sequences of the phyllosphere metagenome *cox* gene homologues which were available on MG-RAST, many of the CODH may be form II types of which the function is not fully known. Therefore, further research into the activity of form II CODH in CO oxidation, along with additional meta-omics investigations of phyllosphere communities, is necessary before the potential of phyllosphere microbiota to oxidise CO is understood.

The phyllosphere CO-oxidising isolate CO17, a member of the *Phyllobacteriaceae*, was most closely related to *Mezorhizobium zhangyense* [87], Some *Mesorhizobium* species are already known to oxidise CO [48], and are common phyllosphere colonisers [88]. However, isolate CO17 and *Mesorhizobium zhangyense* do not cluster with other *Mesorhizobium* species (Fig. 5A). CO17 was also identified as an ‘unclassified Phyllobacteriaceae’ according to the RDP classifier [89] whereas *Mesorhizobium zhangyense* is not currently listed as a validated *Mesorhizobium* species in the LPSN [90], indicating that the taxonomic classification of both of these species requires further attention. Likewise, it is noteworthy that isolate CO17 was isolated from a CO enrichment culture where CO was the sole carbon source and it’s genome contained genes encoding RubisCO (*cbbL* and *cbbS*). Whether this is driving the assimilation of CO2 from CO oxidation remains to be determined; previously identified CO-oxidising *Mesorhizobium* species did not grow on CO as a carbon source.

## Conclusions

Overall, this work demonstrates that the phyllosphere is host to diverse microorganisms capable of oxidising CO. Some of the candidate CO-oxidising species identified here are abundant members of the phyllosphere microbial community and the preliminary functional analyses of phyllosphere communities indicate an average of 25% of bacterial community colonising the phyllosphere contain *cox* (*coxL, coxM* and coxL) gene homologues and therefore may be capable of CO oxidation. Recent research has suggested that carboxydovores capable of CO oxidation at sub-atmospheric concentrations are abundant in both marine and soil samples [59, 60]. As the leaf surface is a nutrient-poor environment with direct exposure to atmospheric CO, we propose that CO oxidation may be a widespread function of the phyllosphere. Further work is required to establish the contribution of phyllosphere microbiota to the global CO cycling and mitigation of this toxic air pollutant.

## List of abbreviations

ANOSIM: Analysis of similarities
CO: Carbon monoxide
CODH: Carbon monoxide dehydrogenase
GC: Gas chromatography
OTU: Operational taxonomic unit
PCR: Polymerase chain reaction
SIMPER: Similarity percentage analysis

## Declarations

### Ethics approval and consent to participate

Not applicable.

### Consent for publication

Not applicable.

### Availability of data and material

The raw 16S rRNA gene sequences from Illumina MiSeq amplicon sequencing were deposited at NCBI Sequence Read Archive (SRA) under the BioProject accession ID PRJNA699530. The 16S rRNA gene sequences of all CO-oxidising candidate species were deposited at NCBI Genbank under the accession numbers MW596379-MW596407. All *coxL* genes from clone libraries were deposited at NCBI Genbank under the accession numbers MW596683-MW596710. The 16S rRNA gene of CO-oxidising isolate CO17 was submitted at NCBI Genbank under the accession MW592845 whereas the assembled genome sequence was submitted under the accession of JAFNIO000000000.

### Author’s contributions

JLP designed the research studies, performed the laboratory experiments, analysed the data and wrote the manuscript. SH assisted with the MiSeq library preparation and analysis of data. EP assisted with data analysis. GDB designed the research studies and edited the manuscript. HS conceived the project, designed the research studies and edited the manuscript. All authors read and approved the final manuscript.

## Acknowledgements

Genome sequencing was provided by MicrobesNG (http://www.microbesng.uk). We would like to thank Dr. Eileen Kröber (Max Plank Institute for Marine Microbiology) for her experimental support.

## Supplementary information

**Additional file 1: Figure S1. Diversity of *coxL* genes used in *coxL* gene primer design.** The tree was deduced using the Neighbour-Joining algorithm implemented in ARB with PAM distance correction based on an alignment of nucleotide sequences [65]. Asterix (*) indicates sequences which were not used in *coxL* primer design.

**Additional file 2: Table S1. Modification of form I *coxL* gene primers**. Mismatches identified in bases are indicated in red. Modifications made to bases in new primer pair are indicated in green.

**Additional file 3: Figure S2. Degradation of CO as a sole carbon source by holly and hawthorn leaf wash communities from heterotrophic enrichment cultures.** Enrichment culture medium contained minimal media with 800_ppm CO and no additional carbon source. A) Woodland hawthorn (HtW) samples. B) Roadside hawthorn (HtR) samples. C) Woodland holly (HlW) samples. Spikes in CO concentration indicate where CO was added to the enrichment culture upon degradation to beyond the detectable limit (<50_ppm).

**Additional file 4: Figure S3. Relative abundance of OTUs enriched in leaf wash cultures containing 800 ppm CO plus yeast extract compared to control cultures without CO.** Samples shown in this figure represent leaf wash cultures which degraded 800 ppm CO, with yeast extract, four consecutive times during three sub-cultures. OTUs which contributed to <2% of total reads on average in the CO enrichment group were placed in the ‘Low abundance’ category.

**Additional file 5: Figure S4. Relative abundance of OTUs enriched with 800 ppm CO as a sole carbon source compared to control cultures without CO.** Samples shown in this figure are those which successfully oxidised four spikes of 800 ppm CO as a sole carbon source after being sub-cultured from CO cultures which contained yeast extract as an additional carbon source. OTUs which contributed to <2% of total reads on average in the CO enrichment group were placed in the ‘Low abundance’ category.

**Additional file 6: Figure S5. Bacterial OTUs identified as indicator species in leaf wash cultures enriched with 800 ppm CO with or without an additional carbon source. A)** Leaf wash OTUs significantly enriched by CO in cultures containing 800 ppm CO plus yeast extract. **B)** OTUs from heterotrophic CO enrichment cultures which were enriched when sub-cultured into cultures containing 800 ppm CO as a sole carbon source. OTUs in red indicate those with P <0.05 and Log_2_ fold change >2. OTUs in blue indicate those with P <0.05 and Log_2_ fold change = <2. OTUs in grey indicate those with P >0.05 and Log_2_ fold change <2. OTUs in green indicate those with P >0.05 and Log_2_ fold change >2.

**Additional file 7: Figure S6. Phylogeny of the RubisCO protein of CO-oxidising isolate CO17.** Bona fide form I Rubisco amino acid sequences were identified using data from Nanba et al., [91]. Phylogeny of CO17 RubisCO and other form I RubisCO sequences was inferred using the Neighbor-Joining method [1] of amino acid sequences in MEGA7 [2]. The percentage of replicate trees in which the associated taxa clustered together in the bootstrap test (500 replicates) are shown next to the branches. The scale bar denotes the number of amino acid differences per site.

