## Supplementary figures and images for "Tree phyllospheres are a habitat for diverse populations of CO-oxidising bacteria"

### Additional file 1

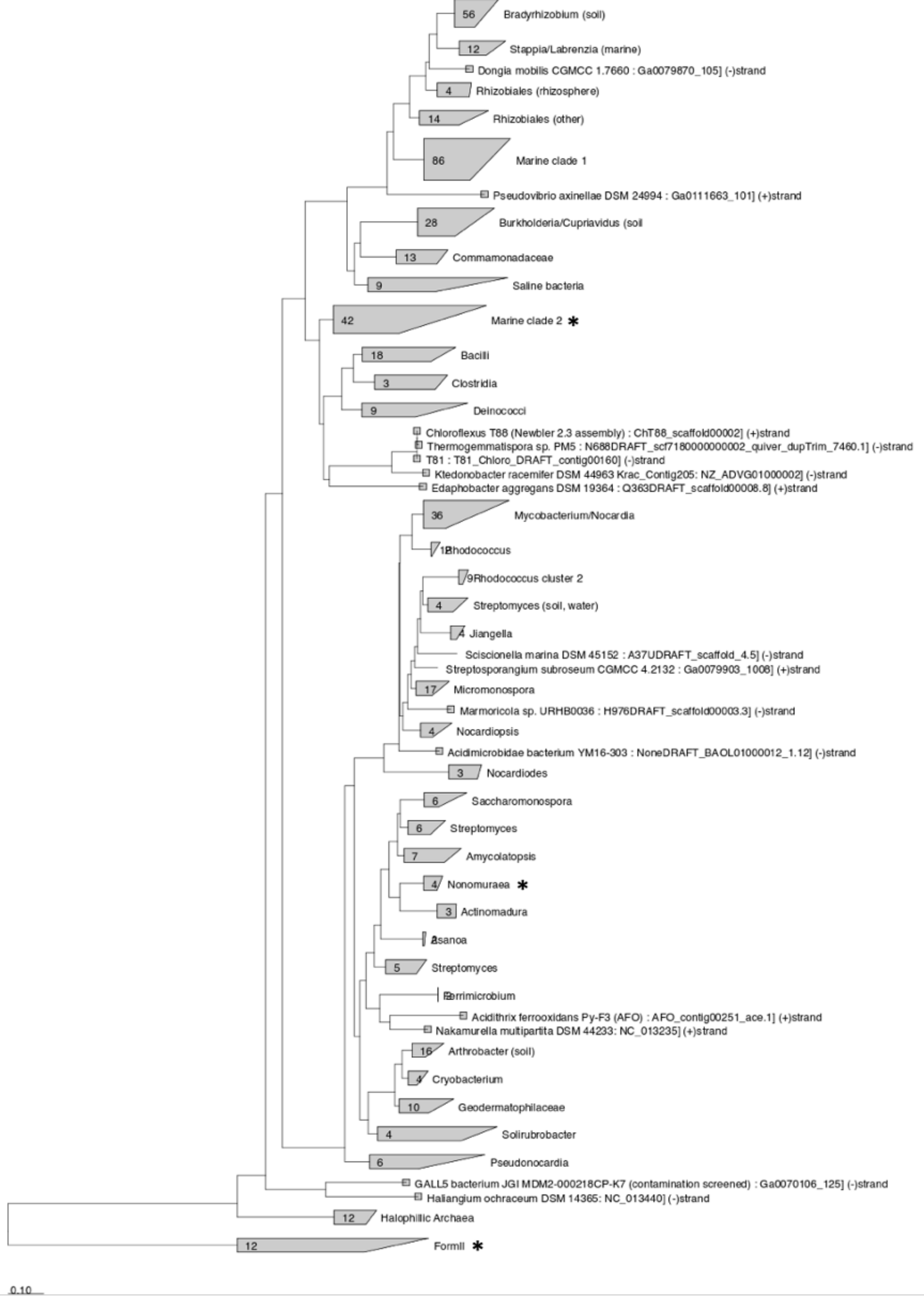

### Additional file 3

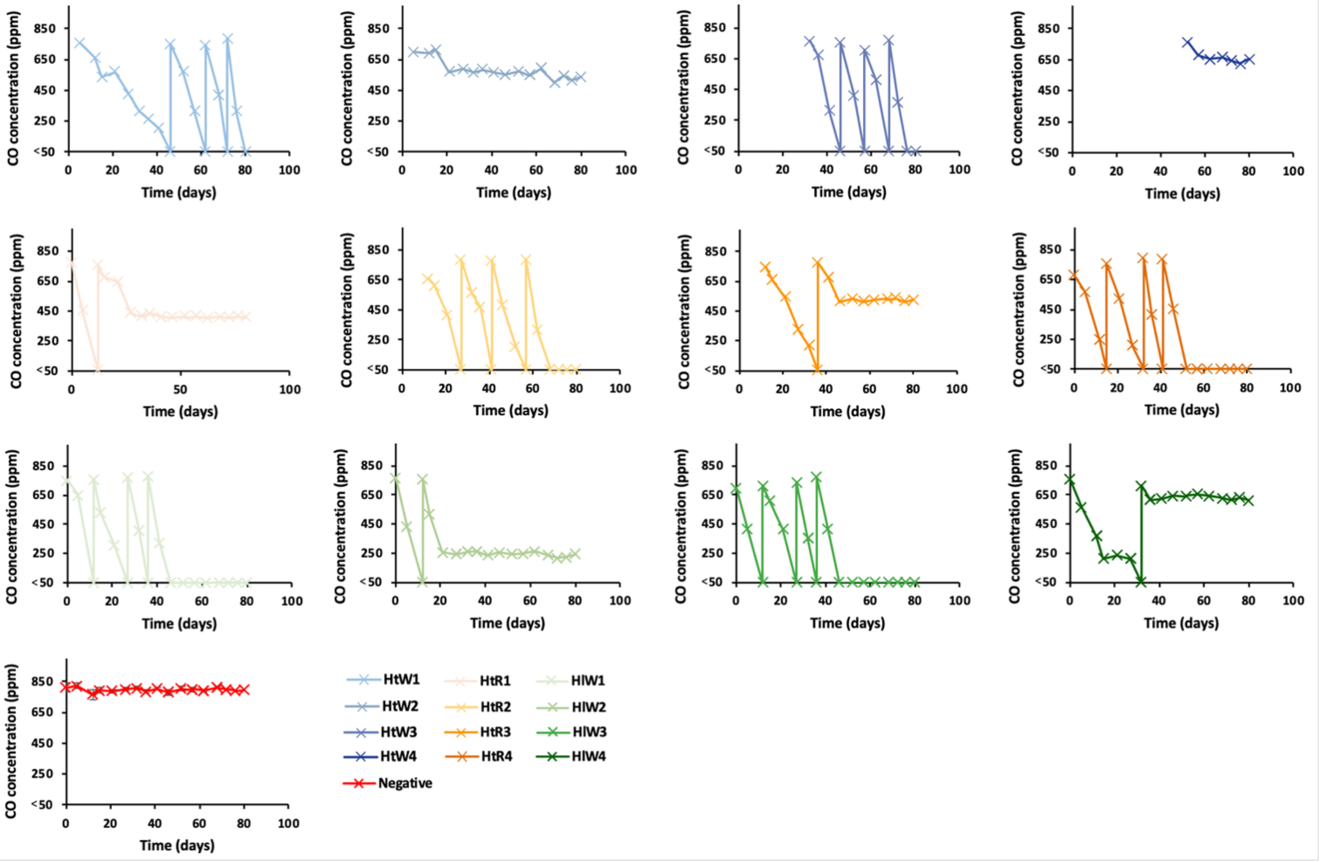

### Additional file 4

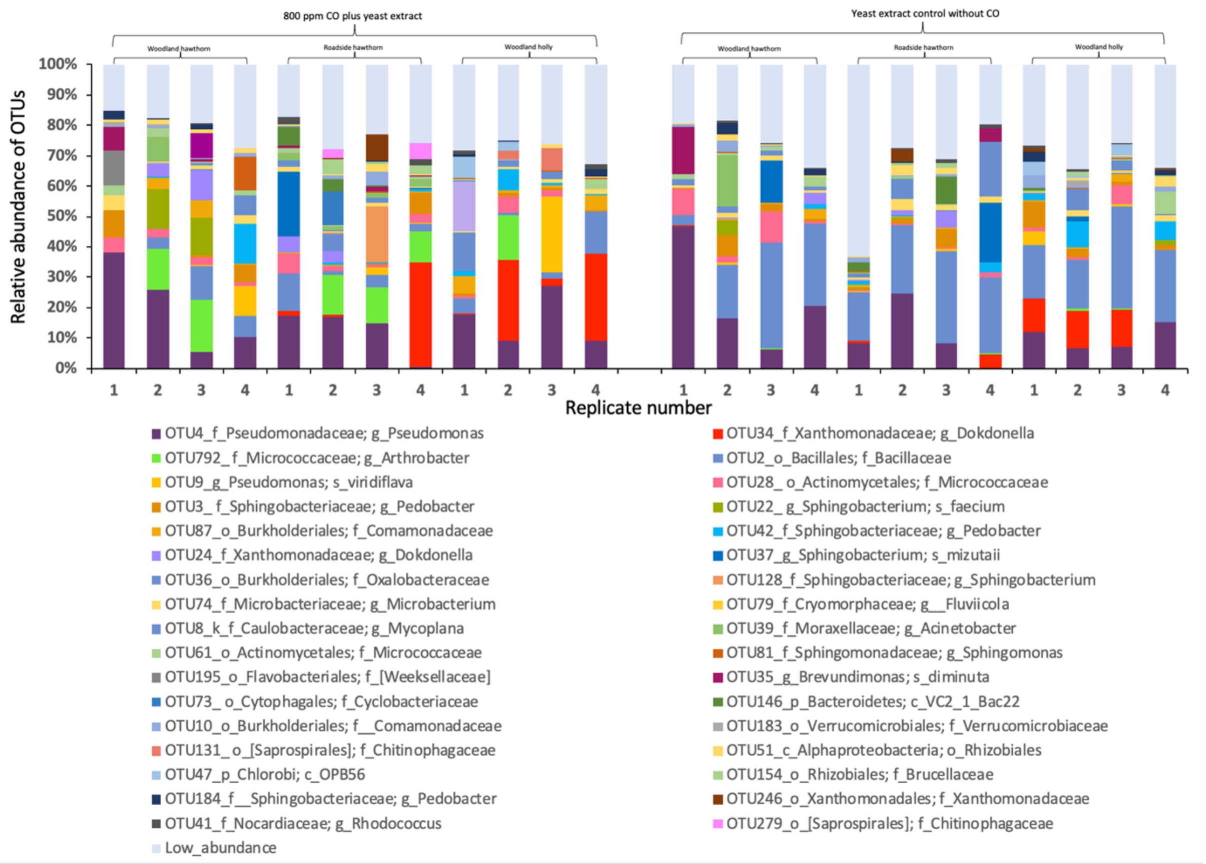

### Additional file 5

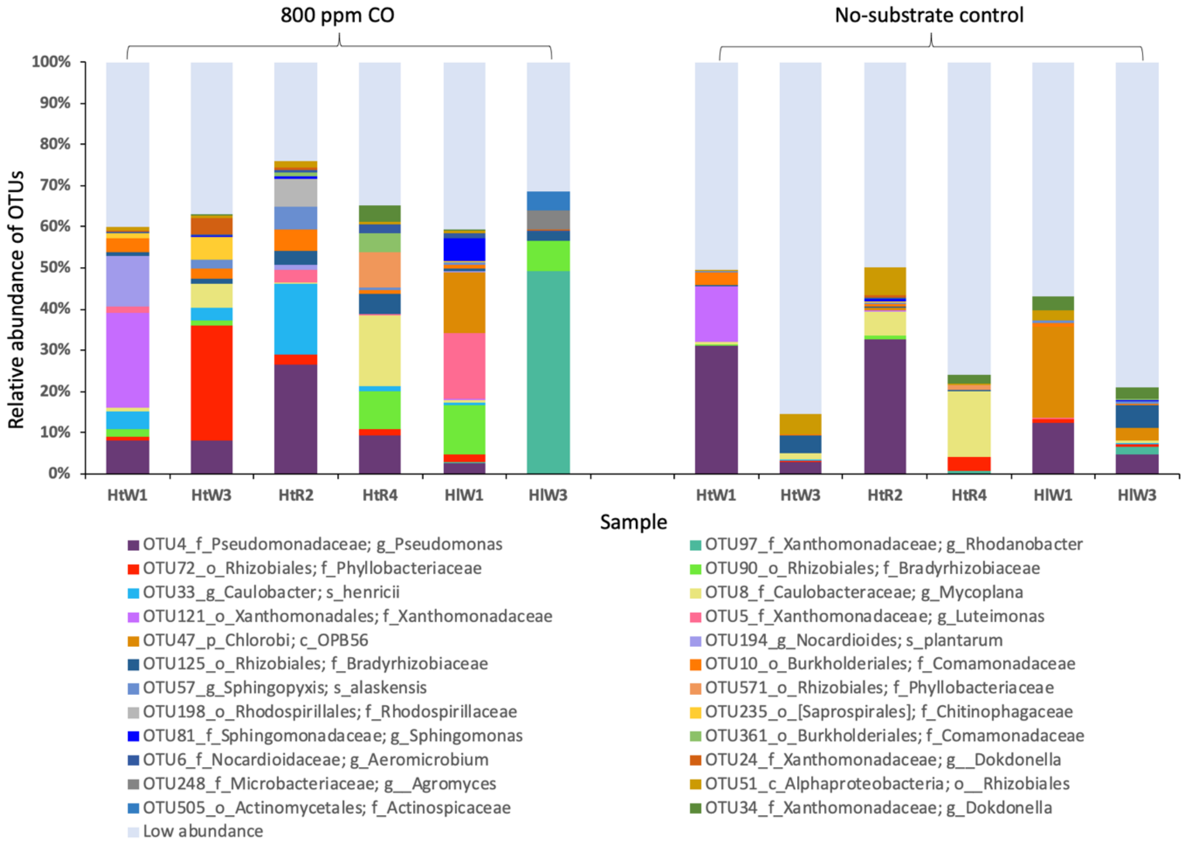

### Additional file 6

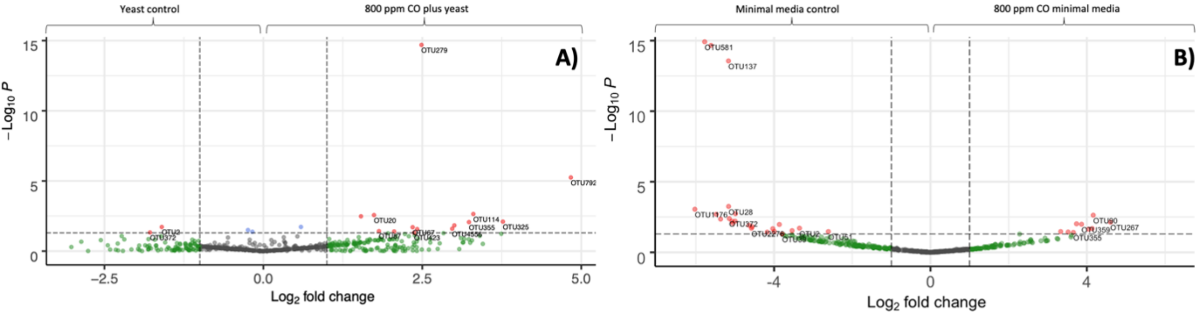

### Additional file 7

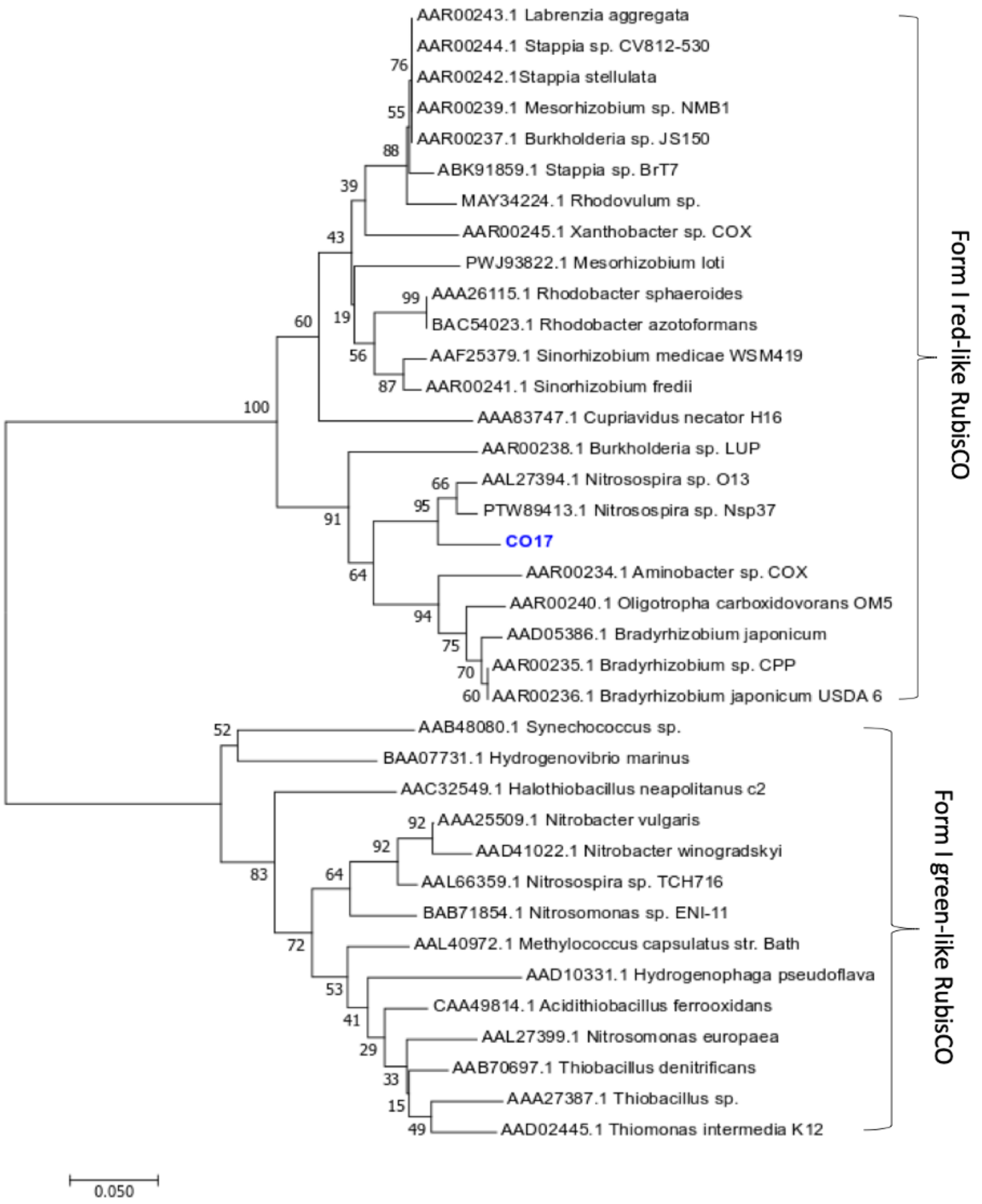
