## Additional file 2 for "Tree phyllospheres are a habitat for diverse populations of CO-oxidising bacteria"

| Primer name | Primer sequence (5’-3’) | Reference |
|---|---|---|
| OMPf | GGC GGC TTY GGS AAS AAG GT | (King 2003) |
| O/Br | YTC GAY GAT CAT CGG RTT GA | (King 2003) |
| Mod_coxL_f | GGC GGN TTY GGN AAY AAR GT | (This study) |
| Mod_coxL_r | YTC DAT DAT CAT NGG RTT DAT | (This study) |
